# Optimized protocols for RNA interference in *Macrostomum lignano*

**DOI:** 10.1101/2023.11.09.565441

**Authors:** Stijn Mouton, Alexandra Mougel, Kirill Ustyantsev, Colette Dissous, Oleg Melnyk, Eugene Berezikov, Jérôme Vicogne

**Affiliations:** European Research Institute for the Biology of Ageing, University of Groningen, University Medical Center Groningen, Groningen 9700AD, The Netherlands; Univ. Lille, CNRS, INSERM, CHU Lille, Institut Pasteur de Lille, U1019-UMR9017 - CIIL - Center for Infection and Immunity of Lille, F-59000 Lille, France

**Author notes:** These authors equally contributed to this work (Stijn Mouton, Alexandra Mougel). Co-corresponding authors., **Authors for correspondence:** Jérôme Vicogne; Stijn Mouton. **Author Contributions**. S.M. wrote the original draft, reviewed and edited the manuscript, realized the project’s conceptualization, contributed to validation, methodology, supervision, funding, and project administration, and provided resources. A.M. wrote the original draft, realized all investigations and visualizations, and performed data curation, validation, and methodology. K.U. wrote the original draft, implemented the dsRNA bacterial extraction method, and performed validation. C.D., O.M., and E.B. reviewed and edited the manuscript, contributed to conceptualization, methodology, supervision, funding, and project administration, and provided resources. J.V. wrote the original draft, reviewed and edited the manuscript, realized the project’s conceptualization, contributed to validation, methodology, supervision, funding, and project administration, and provided resources. All authors have read and agreed to the published version of the manuscript. **Conflict of Interest**. The authors declare no conflict of interest. The funders had no role in the design of the study, in the collection, analyses, or interpretation of data, in the writing of the manuscript, or in the decision to publish the results.

**Keywords:** *Macrostomum lignano*, RNA interference, Electroporation, Soaking, Regeneration, Germline

## Abstract

*Macrostomum lignano*, a marine free-living flatworm, has emerged as a potent invertebrate model in developmental biology for studying stem cells, germline, and regeneration processes. In recent years, many tools have been developed to manipulate this worm and to facilitate genetic modification. RNA interference is currently the most accessible and direct technique to investigate gene functions. It is obtained by soaking worms in artificial seawater containing dsRNA targeting the gene of interest. Although easy to perform, the original protocol calls for daily exchange of dsRNA solutions, usually until phenotypes are observed, which is both time- and cost-consuming. In this work, we have evaluated alternative dsRNA delivery techniques, such as electroporation and osmotic shock, to facilitate the experiments with improved time and cost efficiency. During our investigation to optimize RNAi, we demonstrated that, in the absence of diatoms, regular single soaking in artificial seawater containing dsRNA directly produced in bacteria or synthesized *in vitro* is, in most cases, sufficient to induce a potent gene knockdown for several days with a single soaking step. Therefore, this new and highly simplified method allows a very significant reduction of dsRNA consumption and lab work. In addition, it enables performing experiments on a larger number of worms at minimal cost.

## Introduction

*Macrostomum lignano* is a free-living flatworm that is being developed as a versatile model system for research on a broad range of biological questions (Wudarski *et al*. 2020), including regeneration (Egger *et al*. 2006), *in vivo* stem cell biology (Ladurner *et al*. 2000; Grudniewska *et al*. 2016), germline biology (Pfister *et al*. 2008; Sekii *et al*. 2009; Grudniewska *et al*. 2018), bio-adhesion (Lengerer *et al*. 2014; Lengerer *et al*. 2016), the evolution of sexual reproduction (VIZOSO AND SCHARER 2007; Ramm *et al*. 2019), chromosome evolution (Zadesenets *et al*. 2016; Zadesenets *et al*. 2017), and ageing (Mouton *et al*. 2018). This versatility stems from a convenient combination of biological features of *M. lignano*, which include a transparent body, hermaphroditic reproduction (SCHARER AND LADURNER 2003), short developmental time, and the ability to regenerate the complete body, including the germline. The high regenerative capacity is driven by a pluripotent population of proliferating stem cells called neoblasts (Ladurner *et al*. 2000). In addition, *M. lignano* has significant experimental power thanks to the development of diverse molecular tools and resources. The one feature setting it apart from other flatworm models is the availability of successful methods for transgenesis (Wudarski *et al*. 2017; Ustyantsev *et al*. 2021). In recent years, high-quality genome and transcriptome assemblies (Wasik *et al*. 2015; Grudniewska *et al*. 2016; Wudarski *et al*. 2017; Grudniewska *et al*. 2018), and the transcriptional profiling of the germline and somatic neoblasts have been published (Grudniewska *et al*. 2016). These resources facilitate functional studies and screens using methods such as *in situ* hybridization (ISH) and RNA interference (RNAi). The current commonly used ISH and RNAi protocols in *Macrostomum* were established over a decade ago (Pfister *et al*. 2007; Pfister *et al*. 2008) and are not necessarily optimal.

In *M. lignano*, RNAi is a straightforward method since, like in many other worms (Fontenla *et al*. 2021), they can directly soak up dsRNA dissolved in the medium, probably by an active and efficient mechanism similar to that identified in *C. elegans* (Winston *et al*. 2007; SHIH AND HUNTER 2011; Mcewan *et al*. 2012). To ensure a constant supply of dsRNA to the animals, RNAi studies have always been performed by daily replacing the solution of dsRNA in a culture medium, in which the worms and algae (their food) are maintained. This approach has resulted in the identification of knockdown phenotypes for multiple genes involved in different biological processes, including regeneration, gametogenesis, and adhesion (Sekii *et al*. 2009; Kuales *et al*. 2011; Grudniewska *et al*. 2016; Grudniewska *et al*. 2018; Mouton *et al*. 2021). However, this approach has some significant limitations. The need to daily refresh the dsRNA makes it labour-intensive for larger screens. It also requires a significant amount of dsRNA, which increases the cost of experiments. Moreover, there has never been a clear insight into the efficiency of this approach, raising questions about missing phenotypes during RNAi screens. Therefore, we aimed to develop more convenient, cheaper, highly efficient, and broadly accessible RNAi approaches for *M. lignano*.

Inspired by a recent *C. elegans* publication (Khodakova *et al*. 2021) and our expertise with RNAi in the parasitic flatworm *Schistosoma mansoni* (KRAUTZ-PETERSON *et al*. 2010; Vanderstraete *et al*. 2014), we started these studies by developing an electroporation protocol adapted to this marine organism, with the goal of delivering dsRNA in a single step and resulting in a long-lasting RNA interference. While RNAi by electroporation worked well, during the optimization process, we discovered that it is not the method of delivery of dsRNA that is limiting. Instead, the presence of diatoms in the culture medium is the main parameter that limits the efficient uptake and subsequent efficacy of dsRNA when using the classical soaking protocol.

In this paper, we introduce a simple yet efficient RNAi method for *M. lignano*, based on a single-soak in culture medium with dsRNA. This approach represents an optimal starting point for performing RNAi screens. Moreover, we describe methods to sensitize RNAi, e.g. by electroporation and desalting, to study ‘challenging’ genes. Finally, we introduce a cheap method for the production and purification of bacteria-derived dsRNA, which can be used for *M. lignano* and other model systems.

## Materials and Methods

### Culture of *Macrostomum lignano* and definition of transgenic lines

*M. lignano* is cultured in the laboratory in Petri dishes filled with 32‰ f/2 medium (Andersen 2005) and the diatom *Nitzschia curvilineata* (Bacillariophyceae), which serves as food for the worms. The dishes with diatoms and worms are incubated at 20°C, 25°C or 28°C, 60% humidity, and a 13/11 h light/dark cycle. Worms are weekly transferred to Petri dishes with fresh medium and diatoms and sorted according to developmental stages (eggs, juveniles, adults) to keep track of worm age.

All used transgenic *M. lignano* lines are defined and published in Wudarski *et al*. 2017, and are cultivated in the same way as wild-type animals. To prevent drift in the population, transgenic worms are regularly checked to ensure the expression of fluorescent proteins and eliminate non-fluorescent worms. The NL1 line ubiquitously expresses eGFP under the elongation factor 1 alpha promoter. The NL24 line expresses a single construct with mNeonGreen under the testis-specific promoter of the *Mlig-elav4* gene and mScarlet-I under the ovary-specific promoter of the *Mlig-cabp7* gene.

### RNA extraction and cDNA production

Depending on the conditions tested, worms were not starved or starved in a clean Petri dish containing fresh f/2 medium for 24 h and washed from diatom remains before RNA extraction. Total RNA was extracted from wild-type worms using the Nucleospin RNA® kit (Macherey-Nagel – 740955) or Trizol Reagent™ (Ambion®, #15596018) according to the manufacturer’s protocols, with a few modifications. Briefly, 30 to 80 worms were transferred into a 1.5 mL tube in f/2 medium and placed on ice for a few minutes. f/2 medium was carefully removed as much as possible. Next, 350 µL of RA1 buffer supplemented with 3.5 µL β−mercaptoethanol or 700 µL of TRizol reagent™ were immediately added to the worm pellet. The mixture was vortexed to disrupt worms and eventually stored at -80°C until RNA extraction. RNA was extracted according to the manufacturer’s protocols for the Nucleospin kit and Trizol Reagent but eluted in 22 µL of milliQ water or resuspended in 100 µL, respectively. Next, the RNA was treated with TurboDnase I (Ambion®, AM1907) for 20 min at 37°C to digest traces of genomic DNA.

The AffinityScript Multitemp cDNA Synthesis® kit (Agilent^©^, #200436) was used to convert polyA-mRNA to single-strand cDNA following the manufacturer’s instructions. Briefly, 15.4 µL of total RNA (maximum allowed volume) was used as a template and annealed to the oligo-dT (21) primer. The cDNA synthesis reaction mixture was added, and extension was performed at 50°C for 1 h.

### Enzymatic *in vitro* synthesis of dsRNA

Design and synthesis of dsRNA for the target genes were done as described in Grudniewska *et al*. 2016. Regions of the target genes were PCR amplified from cDNA, genomic DNA, or plasmids with specific primer pairs for each gene (Table S1). The resulting fragments were cloned into the Topo PCR4-TA cloning plasmid (Invitrogen®) or pGEM-T vector system (Promega®), and sequences were validated by Sanger sequencing. The plasmids were used as templates for a new PCR with specific primer pairs extended with a T7 promoter sequence. The quality and specificity of the amplified DNA fragments containing T7 promoters in opposite orientations were assessed by agarose gel electrophoresis. The various dsRNA were next synthesized and purified using the MegaScript RNA kit (Invitrogen®, AM1626M) according to the manufacturer’s protocol. Briefly, 100 to 200 ng of each amplicon were added to the T7 RNA polymerase reaction mixture in a 50 µL final volume and incubated at 37°C for 3 h. Samples were next heated to 75°C in a heat block and slowly cooled down to room temperature (10 to 15 min). This step is critical to promote proper and stable dsRNA annealing. The quality, size, and proper annealing of dsRNA were next validated by agarose gel electrophoresis before DNase I and RNase H treatment. Digested samples were next column-purified, and dsRNA concentration was measured on a Nanodrop spectrophotometer (RNA mode). The integrity of dsRNA was confirmed by electrophoresis, and samples were aliquoted and stored at -20°C, where they were stable for at least 12 months. Yields obtained were usually between 50 to 200 µg of purified dsRNA per reaction.

### Production and purification of bacteria-derived dsRNA

An alternative to the enzymatic *in vitro* synthesis of dsRNA is the production and purification of dsRNA directly from bacteria (Nwokeoji *et al*. 2016; Sturm *et al*. 2018). A detailed step-by-step protocol is provided in Supplementary Materials.

Briefly, exonic fragments of the target genes were cloned into the T444T plasmid vector containing two opposing T7 promoters following T7 terminator sequences flanking the insertion site (Sturm *et al*. 2018). The T444T vector was cut by KpnI and BglII restriction enzymes (NEB) to remove the multi-cloning site insert sequence as close as possible to the T7 promoters. The plasmid backbone was gel-purified from a 1% agarose gel using the FavorPrep GEL/PCR Extraction kit (Favorgen®). Exonic fragments of the target genes were PCR amplified from *M. lignano* gDNA or cDNA using primers containing universal 5’ overhangs to the 5’ and 3’ ends of the linearized T444T backbone (Table S1). The fragments were gel-purified by the freeze & squeeze method, mixed 1:1 with the linearized T444T backbone, and joined using customized low-volume reactions setup by the NEBuilder® HiFi DNA Assembly Master Mix (NEB). The reactions were transformed into the chemically competent HT115 (DE3) *E. coli* strain containing a mutation in the RNAse III gene, insertion of the lactose/IPTG inducible T7 polymerase-encoding gene, and tetracycline resistance gene (Papic *et al*. 2018). Colonies were selected on LB-agar plates supplemented with ampicillin and tetracycline (LB+Amp+Tet), and the presence and length of insertion fragments were checked by colony PCR with a universal T7 promoter primer, followed by 1% agarose gel electrophoresis. Colonies with verified insertions were grown overnight in LB with ampicillin and tetracycline. On the next day, the overnight cultures were used both to create glycerol stocks for long-term storage at -80°C and for plasmid DNA extractions using the Plasmid DNA Extraction Mini Kit (Favorgen®). The extracted plasmids were used for Sanger sequencing using a universal T444T forward sequencing primer (Table S1).

To produce dsRNA, fresh liquid 4 mL night cultures with LB+Amp+Tet were initiated from the prepared glycerol stocks. The next day, 0.2 mL of the starters were inoculated to 20 mL of LB+Amp and allowed to grow for 3 h at 37°C with shaking. Next, 0.1 mL of 0.1 M IPTG was added to initiate the T7 polymerase transcription, and the cells were allowed to grow further for 4 h at 37°C with shaking. The cells were collected by centrifugation, the medium was removed, and the pellets were put at -20°C until further use.

DsRNA species were purified using a modified RNASwift method (Nwokeoji *et al*. 2016). In short, the pellets were resuspended and lysed in 4% Sodium dodecyl sulfate (SDS), 0.5 M NaCl in RNAse-free H_2_O for 10 min at 80°C. Then, proteins, polysaccharides, and SDS were precipitated by adding 0.5 volume of 5 M NaCl and centrifugation. Nucleic acids were then precipitated by adding 1 volume of isopropanol to the supernatants and centrifugation. The pellets were washed two times in 70% ethanol to remove salts and SDS residues, air-dried, and dissolved in 1x DNase I buffer. The samples were then treated by DNAse I (ThermoFisher Scientific®, 18047-019) and RNAse T1 (ThermoFisher Scientific®, EN0541) for 30-60 min at 37°C to remove bacterial genomic DNA, plasmid DNA, and single-stranded RNA molecules from the solutions. To remove low molecular weight non-digestible dsRNA of tRNA and bacterial 5S RNA molecules, any potential SDS residuals, and other contaminants, an additional step of Polyethylene Glycol (PEG8000) assisted precipitation was performed. Briefly, nucleic acids above ∼250 bp in DNA size are precipitated by centrifugation in the presence of 11% PEG and 0.5 M NaCl, while smaller molecules remain in the supernatant. The pellets were washed twice in 70% ethanol, air-dried, and resuspended in RNAse-free water.

The integrity and size of the purified dsRNA were analyzed by 1% agarose gel electrophoresis with ethidium bromide. Rough estimations of the dsRNA concentrations were done by gel-based densitometry, comparing the intensities of the major dsRNA bands to known amounts of 1 kb DNA ladder marker (NEB) bands using Fiji (Schindelin *et al*. 2012). Finally, the samples were diluted to the same concentrations (∼200 ng/µL) and aliquoted. The aliquots were stored at -20°C for several months.

### dsRNA electroporation

After starving for 24-48 h, worms were transferred into a 12-well plate containing diluted f/2 medium (4df/2, 1/4 dilution of f/2 in ddH2O) for 1 h at 20°C. Worms were then transferred to electroporation cuvettes (4 mm Gap – FisherBrand - #FB104) containing 400 µL 4df/2 and 2 µg dsRNA. After gentle agitation to detach worms, an electrical pulse (single pulse square signal at 60 V for 20 ms) was immediately performed in a Gene pulser Xcell – CE module (Biorad®). Worms were left to rest for 2 h in the cuvette at room temperature, then transferred into a 12-well plate containing 1 mL 4df/2, and dead/damaged worms were eliminated. The plate with worms was incubated overnight at 20°C. The next day, 500 µL of f/2 (undiluted) was added, and this was repeated 3-4 h later to resalt progressively. Finally, 3-4 h later, worms were transferred to a 12-well plate containing f/2 with diatoms and maintained at 20°C. At different times, the worms were then cut, imaged, or harvested depending on the experimental conditions. Harvested worms were pelleted and stored at -80°C.

### dsRNA soaking

After starving for 24 - 48 h, worms were transferred into a 12-well plate in 400 µL of f/2 or 4df/2 medium, and 2 µg of dsRNA was added. After 24 h at 20°C, the medium was replaced by 1 mL of f/2 containing diatoms. When using 4df/2, after 24 h, 500 µL of f/2 (undiluted) was added, and this was repeated 3-4 h later. Finally, the worms were transferred to a 12-well plate containing f/2 with diatoms. After treatments, worms were maintained at 20°C or 28°C depending on experiments. Then, they were cut, imaged, or harvested at a given time, depending on the experimental conditions. Harvested worms were pelleted and stored at -80°C.

### qPCR quantification

For quantitative PCR (qPCR), cDNA was amplified using the Brilliant III ultra-fast SYBR Green QPCR Master Mix® (Agilent^©^, #600882) in a 20 µL final volume in 96-well PCR optical plates (Axygen`®`). Primers were used at 250 nM final concentration (see primer sequences in Table S1) and the equivalent of 1 µL of cDNA per reaction. qPCR was performed using the QuantStudio 3 detection system (Applied Biosystems^®^) with the following steps: initial denaturation 95°C for 5 min, denaturation at 95°C for 10 s, annealing/extension at 63°C for 20 s, 40 cycles, and final denaturation step to identify the T_m_ of amplicons. The percentage of residual mRNA was determined following normalization with the S19 ribosomal protein as a housekeeping gene according to the 2^-ΔΔCt^ method.

### Western blot for GFP and fluorescence quantification on the NL1 strain

After dsRNA electroporation or soaking, 30 worms (NL1 strain or NL10 as reference) were pelleted and lysed with 100 μL lysis buffer (20 mM HEPES, 1% NP40, 0.1% SDS, 5% glycerol, 142 mM KCl, 5 mM MgCl_2_, 1 mM EDTA, pH 7.45) supplemented with phosphatase inhibitors (1/200 Phosphatase Inhibitor Cocktail 2 – Sigma® P5726) and protease inhibitors (1/400 Protease Inhibitor Cocktail – Sigma® P8340) for 20 min on ice. Lysates were centrifuged (20 000 x G at 4°C), and the supernatant was collected. The total protein concentration was determined with the BCA Protein Assay Reagent Kit (Pierce® *#*10741395), and equal amounts of proteins (∼20 µg) were resolved on NuPAGE 4–12% Bis-Tris gels (Novex® by Invitrogen™ - WG1401BX10) under reducing conditions. Separated proteins were transferred onto a polyvinyl difluoride (PVDF) membrane in Towbin buffer (10% methanol, 10% Tris-glycine 1X, 0.0025% SDS). The membrane was then equilibrated in a blocking buffer (8 g casein/L, PBS 1X, 0.2% Tween). Proteins were analyzed by western blotting with GFP antibody (B-2, Santa Cruz Biotechnology® – sc 9996) and GAPDH antibody (Ambion® – AM4300). After incubation with the appropriate species-specific horseradish-peroxidase-conjugated secondary antibodies (anti-mouse #115-035-146 – Jackson ImmunoResearch Lab®) and incubation with West Dura Extended Duration substrate (ThermoScientific® #10445345), the antigen-antibody complexes were captured by digital imaging with a cooled charge-coupled device (CCD) camera (LAS 4000, Fuji, Tokyo, Japan).

GFP fluorescence was measured in black 384-well plates (Greiner bio-one® - #781091) using an Infinite M200 pro reader (Tecan™). Typically, 20 µL of worm lysate was completed with 40 µL of lysis buffer, and measurements were performed using 488 nm and 520 nm as excitation and emission wavelengths respectively. The signal was normalized to the protein content for each sample. Protein concentration was determined using a BCA Protein assay kit (Pierce® *#*10741395) with 5 µL of worm lysate.

### Carmine Red staining and imaging

Worms were relaxed in 7.14% MgCl_2_ in ddH2O for 10 min and fixed in 4% PFA (Thermo Scientific® - #28908) in concentrated PBS (0.1 M Sodium Phosphate pH 7.5, 1.5 M NaCl) for 10 min at room temperature in a Petri dish. Worms were then transferred in 1.5 mL tubes and washed with 50% ethanol, followed by 70% ethanol, for 5 min each. They can be stored at 4°C for weeks. Next, they were stained with Carmine red solution (5 mg Carmine Red (Merck® – C1022), 5 ml 37% HCL, 5 ml water, and after one hour 200 mL 90% ethanol was added; the solution was boiled at 90°C, and an additional 200 ml 90% ethanol was added) for 50 min at room temperature, filtered and stored in the dark. Stained worms were washed with 1 mL of acidic ethanol (HCl 0.9% / 70% ethanol), then with 90% and 100% ethanol. Worms were mounted between a slide and coverslip using warmed (60°C) Balsam (Merck® – # C1795) and cooled down at room temperature. Slides are stable for months and were imaged on a Zeiss LSM 880 using the 488-laser beam and a 520-695 nm band path, and/or the AiryScan2 Module system in SR (super-resolution) mode.

### Amputation and visualization of regeneration

Worms were relaxed in a drop of 7.14% MgCl_2_ in ddH2O on a Petri dish, and the whole body or tail plate was amputated using a fine steel surgical blade (Swann-Morton – SM61). For whole-body amputation, the worms were cut at the anterior tip of the testes. For the tail amputation, worms were cut posterior to the ovaries. After amputation, worms were transferred to fresh f/2 medium and diatoms. Photos of regenerating worms were taken with an EVOS XL Core Imaging System (ThermoFisher Scientific®). For this, worms were individually relaxed in a drop of 7.14% MgCl_2_ in ddH2O on a Petri dish, after which most of this solution was removed to squeeze the worm slightly without damaging it.

### Fluorescent imaging of phenotypes

Fluorescent imaging was performed using a Zeiss Axio Zoom V16 microscope with an HRm digital camera and Zeiss filter sets 38HE (FITC) and 43HE (DsRed). For this, worms were individually relaxed in a drop of 7.14% MgCl_2_ in ddH2O on a Petri dish, after which most of this solution was removed to slightly squeeze the worm without damaging it.

## Results

### Electroporation as a method to knockdown gene expression in *Macrostomum lignano*

The current method to knockdown gene expression in *M. lignano* is based on dsRNA soaking. This interference is performed by maintaining worms in a solution of dsRNA diluted in f/2, an enriched seawater medium. Even though this approach enabled the identification of several gene functions, the number of worms simultaneously treated is limited, and, importantly, it requires daily renewal of dsRNA solution, which is time-consuming, expensive, and requires many worm handlings. To improve this protocol, we adapted the electroporation method used on nematodes (Khodakova *et al*. 2021) and another flatworm, the parasite *Schistosoma mansoni*. Contrary to *S. mansoni* which is electroporated in an isotonic buffer with low salt concentration (PBS or DMEM media) (Neves *et al*. 2005; Vanderstraete *et al*. 2014), *M. lignano* cannot be directly electroporated in f/2 since the high salt concentration (32 g/L) will generate a high current, sparks, and heating in the cuvette resulting in massive damages and worm death. Therefore, it was necessary to first transfer the worms to a lower salt solution and let them adapt. To adjust osmolarity to the level of PBS or other cell culture media, we diluted f/2 four times (around 8 g/L). Worms can easily adapt to this diluted f/2 (4df/2) and then be exposed to the electric pulse (RIVERA-INGRAHAM *et al*. 2013; RIVERA-INGRAHAM *et al*. 2016). However, afterwards, a progressive adjustment to regular f/2 by incremental addition of salt was beneficial, probably due to local alteration of the epidermis. To keep the electroporation medium clean and limit spark risk, worms were starved for 24 h in f/2 to allow them to excrete digested diatoms, and were washed before their transfer into 4df/2. Since electroporation parameters will strongly impact worm damage and mortality, we first determined the appropriate voltage allowing the best survival rate of worms after one day and four days (Fig. 1A). The optimal voltage to recover a maximum of macroscopically undamaged worms (no cysts and dynamic swimming and surface attachment) was 60 V (1 pulse 200 ms, square signal), while for Schistosomes 100 V is classically used (KRAUTZ-PETERSON *et al*. 2010; Vanderstraete *et al*. 2014).

**Figure 1:**
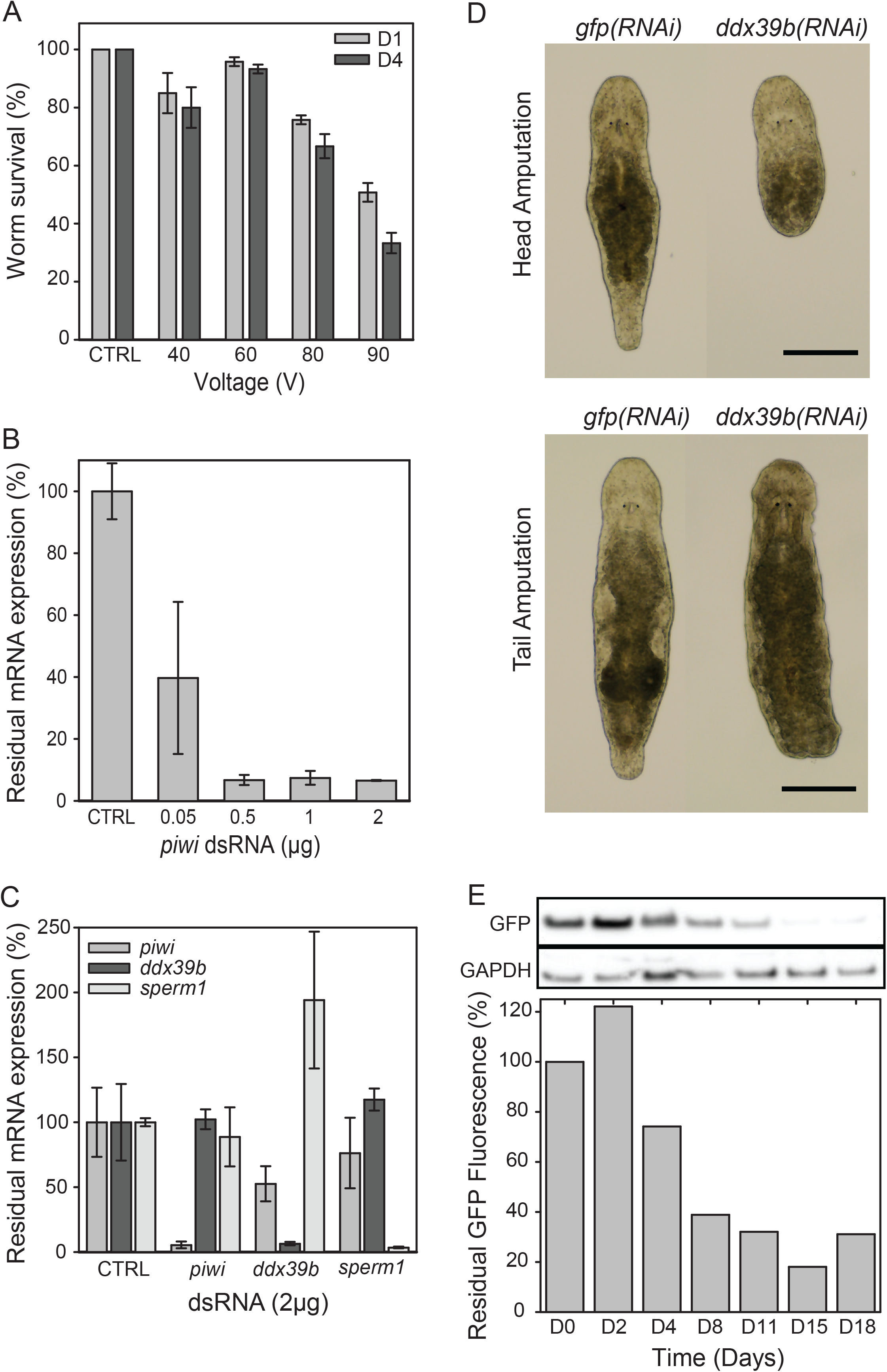
Gene expression interference using electroporation. A. Worm survival rates (percentage of control +/- S.D.) at 1 and 4 days (D1; D4) after electroporation according to the voltage applied (40 worms, n=3, CTRL: no electric pulse). B. Quantification of residual *piwi* mRNA expression using RT-qPCR (percentage residual expression) 4 days after electroporation with different amounts of *piwi* dsRNA per cuvette (µg of dsRNA in 400 µL, 75 worms per assay, n=3, +/- S.D., CTRL: no dsRNA and no electric pulse) C. Quantification of residual *piwi*, *ddx39b* and *sperm1* mRNA expression using RT-qPCR (percentage residual expression) 4 days after electroporation of 2 µg (5 ng/µL) of *piwi*, *ddx39b,* and *sperm1* dsRNA (80 worms per assay, n=3, +/- S.D., CTRL: no dsRNA and no electric pulse). D. Impairment of whole-body and tail regeneration after electroporation with *ddx39B* dsRNA. RNAi was performed by electroporating wild-type worms with 2 µg dsRNA. *gfp* interference represents the negative control. Amputations were performed 3 days after electroporation, and photos were taken 1 week after amputations. For each condition, 15 worms were studied. Scale bars: 200 µm. E. Quantification of residual *gfp* expression after electroporation of 2 µg *gfp* dsRNA (5 ng/µL, 300 worms total). For each time point up to 18 days, 30 worms were selected. GFP fluorescence was measured (Ex=488 nm, Em=520 nm) and normalized to protein content as a percentage of fluorescence at D0. For each time point, residual GFP expression was revealed by Western blot (14 µg total protein load) with an anti-GFP antibody (27 kDa). GAPDH is used as a loading control (36 kDa).

Next, we evaluated the efficacy of RNAi by electroporation using a well-characterized gene, *Mlig-piwi*, an argonaute family nuclease involved in the maintenance of germline, neoblasts, and regeneration in *Macrostomum* (DE MULDER *et al*. 2009; Zhou *et al*. 2015). We first determined the required amount of dsRNA to induce a significant knockdown (Fig 1B). To perform this study quantitatively, we used RT-qPCR to measure the residual *piwi* mRNA copies. Surprisingly, with a dsRNA dose as low as 0.5 µg of dsRNA in 400 µL (∼1.25 ng/µL), we measured a >90% reduction of *piwi* RNA copies after 4 days on a population of 80 worms (n=3), compared to controls without dsRNA.

We extended this analysis to two other genes: *Mlig-sperm1*, involved in spermatogenesis (Grudniewska *et al*. 2018), and *Mlig-ddx39b,* involved in neoblast function and regeneration (Grudniewska *et al*. 2016). We measured the residual level of mRNA 4 days after electroporation with 2 µg of dsRNA (Fig. 1C). For each targeted gene, *piwi*, s*perm1,* or *ddx39b*, the expression was strongly reduced (>90%), confirming the efficacy of RNAi. However, we also observed modifications in the expression levels of their two counterparts. This can be explained by cross-regulations between p*iwi*, *ddx39b,* and s*perm1* genes since they all play a central role in stem cell regulation and gametogenesis. These hyper- or hypo-activations might also be due to potential damages caused by electroporation and the consequent regeneration needs. Even with the compelling RT-qPCR data that confirmed the knockdown of gene expression, we wanted to demonstrate that RNAi by means of electroporation can result in well-characterized regenerative phenotypes (Grudniewska *et al*. 2016). For this, we performed *ddx39b* RNAi experiments, during which worm heads or tails were amputated three days after electroporation (Fig 1D). The regenerative capacity was evaluated one week later at 20°C. For the negative controls, all amputated heads could regenerate a new body, and all worms with an amputated tail could regenerate the tail. For both amputation levels, *ddx39b*(*RNAi*) worms could not regenerate at all, as there was no blastema formation.

To evaluate the duration of the effect of this single electroporation treatment, we preferred to study a non-deleterious phenotype at the protein level to keep a healthy worm population. This was done by performing RNAi against *gfp* using the transgenic NL1 line, which ubiquitously expresses enhanced *gfp* (Wudarski *et al*. 2017). A population of 300 worms was electroporated with 2 µg of *gfp* dsRNA, and the fluorescence intensity of GFP was measured together with Western blot analysis on a subset of 30 worms at frequent time intervals for 18 days (Fig. 1E). Interestingly, this interference against constitutive expression of *gfp* was very efficient but required 8 - 11 days to be fully effective at the protein level and lasted at least up to 15 days when performed at 20°C.

Altogether, these data demonstrate that electroporation is a powerful method to induce RNAi in *M. lignano* on many worms with a single treatment and over a long period. However, when analyzing a very strong phenotype such as *ddx39b(RNAi*), which induces a rapid death of worms by depletion of stem cells (Grudniewska, 2016 #3), we observed a massive loss of worms in the control condition, *i.e.* non-electroporated but containing the *ddx39b* dsRNA, within two weeks. Therefore, we wondered if the low salinity and osmotic stress imposed during the electroporation procedure could be sufficient to make enough dsRNA penetrate into the worms independently of the electric pulse.

### Effect of low salinity on RNAi by soaking in *M. lignano*

To investigate the contribution of osmotic stress to RNAi efficacy, we analyzed the lethal effect of *ddx39b* dsRNA soaking on worms in 4 times diluted f/2 (4df/2). A single group of 30 worms (n=3) was incubated in 400 µL of 4df/2 containing 2 µg of dsRNA in (5 ng/µL) and split into 2 electroporation cuvettes, respectively with or without the electric pulse (Fig. 2A). After progressive “resalting” in standard f/2 for 12 h, and elimination of dead/damaged worms, we measured worm viability for 14 days at 28°C to accelerate the appearance of the lethal phenotype. We also used *gfp* dsRNA, with or without an electric pulse, as an additional negative control. As expected, electroporation of *ddx39b* dsRNA led to a complete death of the worm population within 14 days, but surprisingly, 98% of worms without the electric pulse died as well. The initial rate of death was, however, faster for the electroporated worms, suggesting that the electric pulse damaged the worms and thus required higher regeneration needs. This deleterious effect of electroporation up to 14 days is confirmed by the no dsRNA and *gfp(RNAi)* controls showing a lower survival rate (80 and 60%, respectively) compared to non-electroporated controls (>90%).

**Figure 2:**
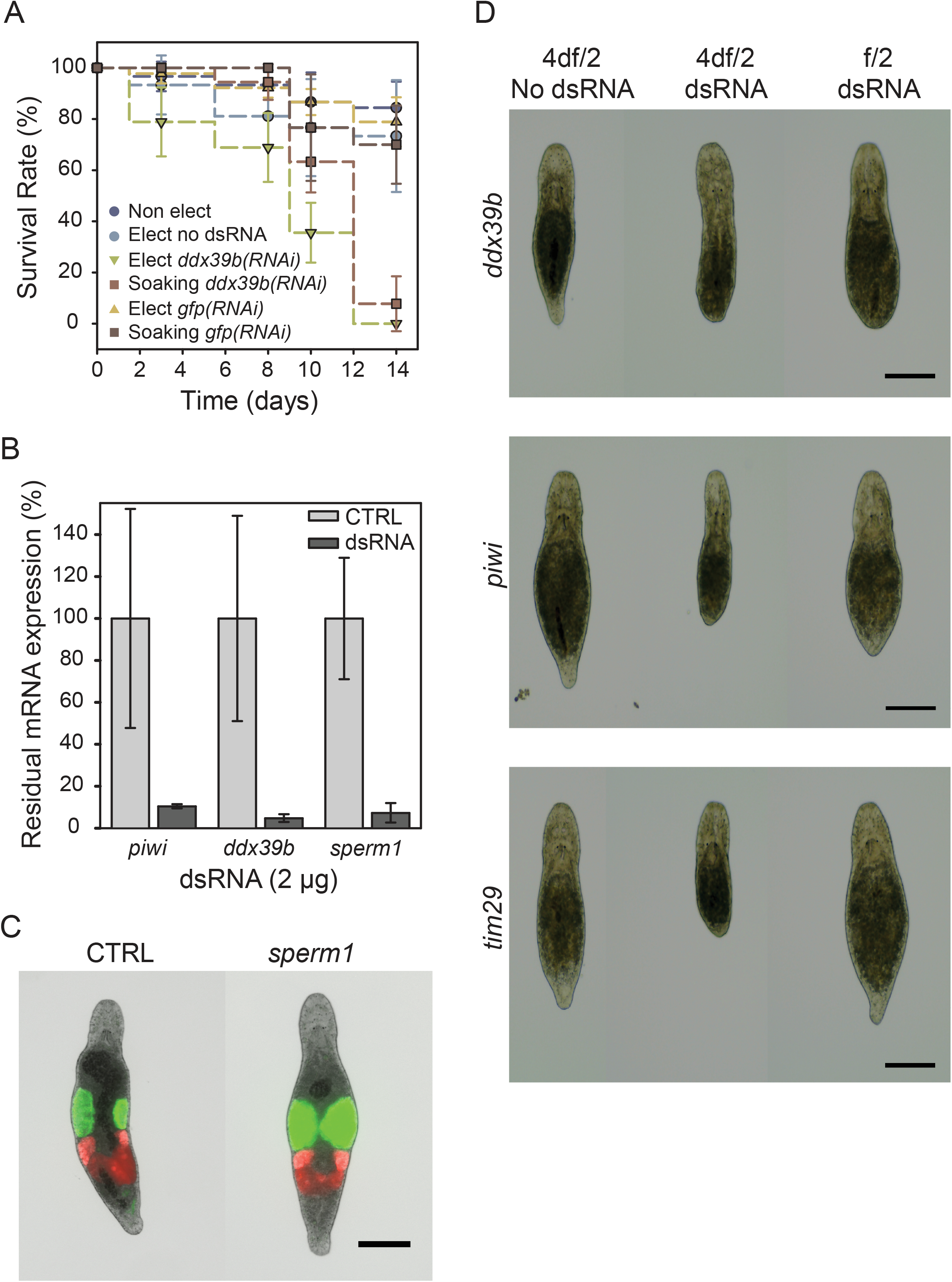
Gene expression interference by single soaking in diluted f/2 (4df/2). A. Worm survival rate (percentage of control) after electroporation or single soaking with 2 µg (5 ng/µL) of *ddx39b* dsRNA in 4df/2 at 28°C for 14 days (30 worms per condition in 400 µL, n=3, +/- S.D.). B. Quantification of residual *piwi*, *ddx39b* and *sperm1* mRNA expression using RT-qPCR (percentage of residual expression, +/- S.D.) 4 days after single soaking with 2 µg of *piwi*, *ddx39b,* and *sperm1* dsRNA in 4df/2 (80 worms per assay, n=2, CTRL: no dsRNA). C. Single soaking in 2 µg *sperm1* dsRNA causes swelling of the testes. Using the NL24 transgenic line, this can be easily visualized due to the testes-specific expression of *mNeonGreen* (15 worms per condition, CTRL: no dsRNA, photos taken 14 days after the single soak, scalebar: 200 µm). D. Regenerative phenotypes after a single soak in *ddx39b*, *piwi*, or *tim29* dsRNA (2 µg). *ddx39b* interference results in a complete lack of regeneration in both 4df/2 and f/2 medium. *Piwi* interference results in impaired regeneration in both 4df/2 and f/2 mediums. *Tim29* interference only impairs regeneration when the single soak is performed in 4df/2. (15 worms per condition, body amputated 5 days after soaking; photos taken 4 days after amputation, scale bars: 200 µm).

Based on these observations and knowing that *ddx39b* knockdown generates a very strong lethal phenotype, we extended these investigations by measuring the remaining expression of *piwi* and s*perm1* by RT-qPCR, in addition to *ddx39b*. Here, we performed a single soaking in 4df/2 for 2 h, followed by progressive salt readjustment at 20°C (Fig. 2B). Surprisingly, after 4 days, we measured knockdown levels >85% and similar to those obtained by electroporation. Altogether, these data suggest that osmotic stress is sufficient to efficiently knock down gene expression by a single soaking in 4df/2 with dsRNA.

Since osmotic stress can promote worm alterations, we analyzed phenotypes obtained with this approach. First, we evaluated the impact of knocking down *sperm1* expression in the NL24 transgenic line, expressing mNeonGreen in the testes and mScarlet-I in the ovaries (Wudarski *et al*. 2017). As published before, in worms exposed to *sperm1* dsRNA, the testes clearly enlarge due to the accumulation of spermatozoa (Fig. 2C). Second, we studied the impact of osmotic stress on driving regenerative phenotypes with *ddx39b*, *piwi*, and *Mlig-tim29* dsRNA (Fig. 2D). *Tim29* was added as it represents a more ‘challenging’ phenotype which is only observed in more extreme conditions such as whole-body regeneration and is very variable between worms (Mouton *et al*. 2021). Besides the negative control that lacks dsRNA during desalting, we also added a single soak in regular f/2 with dsRNA. Osmotic stress during dsRNA exposure without electroporation can clearly block regeneration during *ddx39b*(*RNAi*) and impair regeneration during *piwi*(*RNAi*) and *tim29*(*RNAi*). Surprisingly, exposure to dsRNA during a single soak in a regular f/2 medium also caused an apparent regenerative phenotype for *ddx39*(*RNAi*) and *piwi*(*RNAi*) (Fig. 2D).

Because of the observed regenerative phenotypes during *ddx39b(RNAi)* and *piwi(RNAi)* during a single soak without electroporation or osmotic stress, we wondered if a single soak in regular f/2 with dsRNA could still be sufficient to efficiently knockdown gene expression.

### A single soaking in regular f/2 is sufficient to induce strong RNAi in *Macrostomum lignano*

To further investigate the contribution of osmotic stress on the RNAi efficacy, we chose to repeat the *ddx39b*(*RNAi*) survival assay at 28°C but with regular f/2 in the non-electroporated group (Fig. 3A). As expected, *ddx39*(*RNAi*) using electroporation led to a 100% of death of worms within 14 days. Interestingly, also in the single soaking group, >90% of worms died within the same period of time. This experiment suggests that a single soaking of worms in regular f/2 containing dsRNA is sufficient for the *ddx39b*(*RNAi*) phenotype to appear fully. To verify if this observation could be extended to other targets, we extended experiments to *piwi* and *sperm1* (Fig. 3B). Worms were soaked for 2 h in 400 µL of f/2 containing 2 µg of *piwi*, *ddx39b* or *sperm1* dsRNA. RNA transcription levels were analyzed by RT-qPCR, and the observed knockdown was above 85% for each gene.

**Figure 3:**
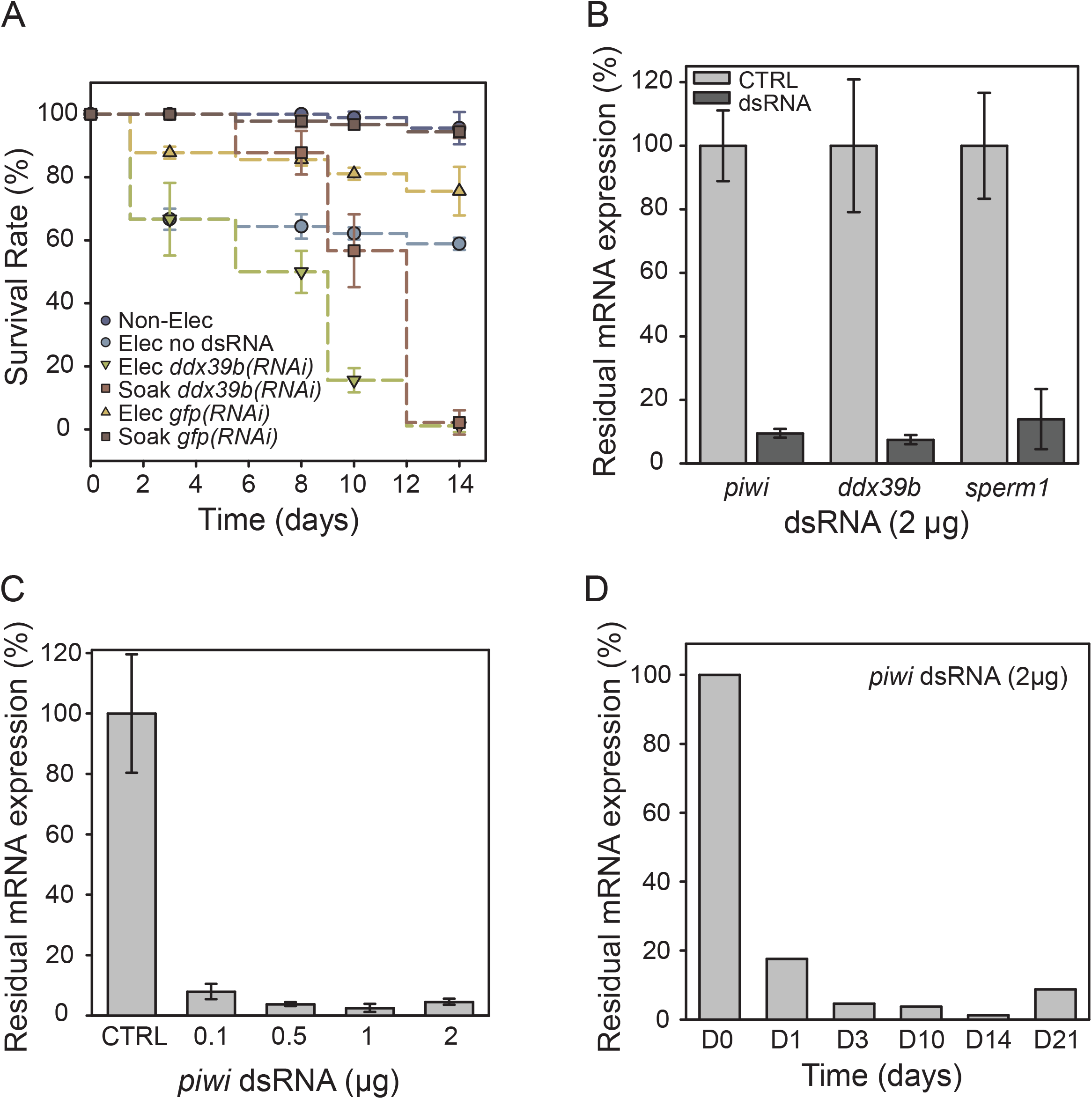
Single soaking in f/2 is sufficient to promote efficient interference. A. Worm survival rate (percentage of control) after electroporation or single soaking in f/2 with 2 µg of *ddx39b* or *gfp* dsRNA (5 ng/µL) at 28°C for 14 days (30 worms per condition in 400 µL, n=3, +/- S.D.). B. Quantification of residual *piwi*, *ddx39b* and *sperm1* mRNA expression using RT-qPCR (percentage of residual expression) 4 days after single soaking with 2 µg (5 ng/µL) of *piwi*, *ddx39b* and *sperm1* dsRNA in f/2 (80 worms per assay in 400 µL, n=2, +/- S.D., CTRL: no dsRNA and no electric pulse). C. Quantification of residual *piwi* mRNA expression using RT-qPCR 4 days after soaking with different concentrations of *piwi* dsRNA (2 µg in 400 µL per wells, 80 worms per assay, n=3, +/- S.D., CRTL: no *piwi* dsRNA). D. Quantification of residual *piwi* mRNA expression using RT-qPCR after single soaking with 2 µg of *piwi* dsRNA (5 ng/µL) up to 21 days (600 worms total, 80 worms analyzed per day).

Next, we determined the amount of dsRNA required to obtain the maximal knockdown at this condition. Worms were single-soaked for 2 h with increasing amounts of *piwi* dsRNA from 0.1 to 2 µg in 400 µL (0.25 to 5 ng/µL). The remaining expression of *piwi* mRNA after 4 days was analyzed by RT-qPCR and confirmed that efficient knockdown is already obtained by a dsRNA concentration as low as 0.25 ng/ µL, *i.e.* 0.1 µg for a population of 80 worms in 400 µL (Fig. 3C).

The traditional RNAi protocol for *M. lignano* recommends renewing the dsRNA soaking at least daily until the expected phenotypes appear (Pfister *et al*. 2008). To investigate if a single soaking is sufficient to induce a sustained knockdown, a group of 600 worms was soaked once in *piwi* dsRNA (6 groups of 100 worms with 2 µg of dsRNA in 400 µL and pooled together after 2 h), and 80 worms were analyzed by RT-qPCR for measurement of *piwi* mRNA levels after 1, 3, 10, 14 and 21 days. Interestingly, the knockdown was observed already after one day (>80%), was strongly sustained for 14 days (>95%), and began to relapse after 21 days (Fig. 3D).

To complement this quantitative analysis, we explored the *sperm1* phenotype by applying the Carmine Red staining protocol to *M. lignano*. This staining is regularly used to image whole-mount *S. mansoni* adult worms, which are not transparent and much thicker than *Macrostomum.* The value of the Carmine Red staining for *M. lignano* is that not only organs but also cellular organization can be easily observed by confocal microscopy (Supplementary Fig. S1), similar to *S. mansoni* (Neves *et al*. 2005; Collins *et al*. 2011). Different cell types can also be identified, which is particularly convenient for interpreting germline-related phenotypes (Supplementary Fig S1; Supplementary Video S1 and S2). These experiments confirmed that a single soak in regular f/2 with dsRNA is sufficient to obtain the typical *sperm1* phenotype, characterized by an accumulation of spermatozoa in the testes, which causes their enlargement (Fig. 4).

**Figure 4:**
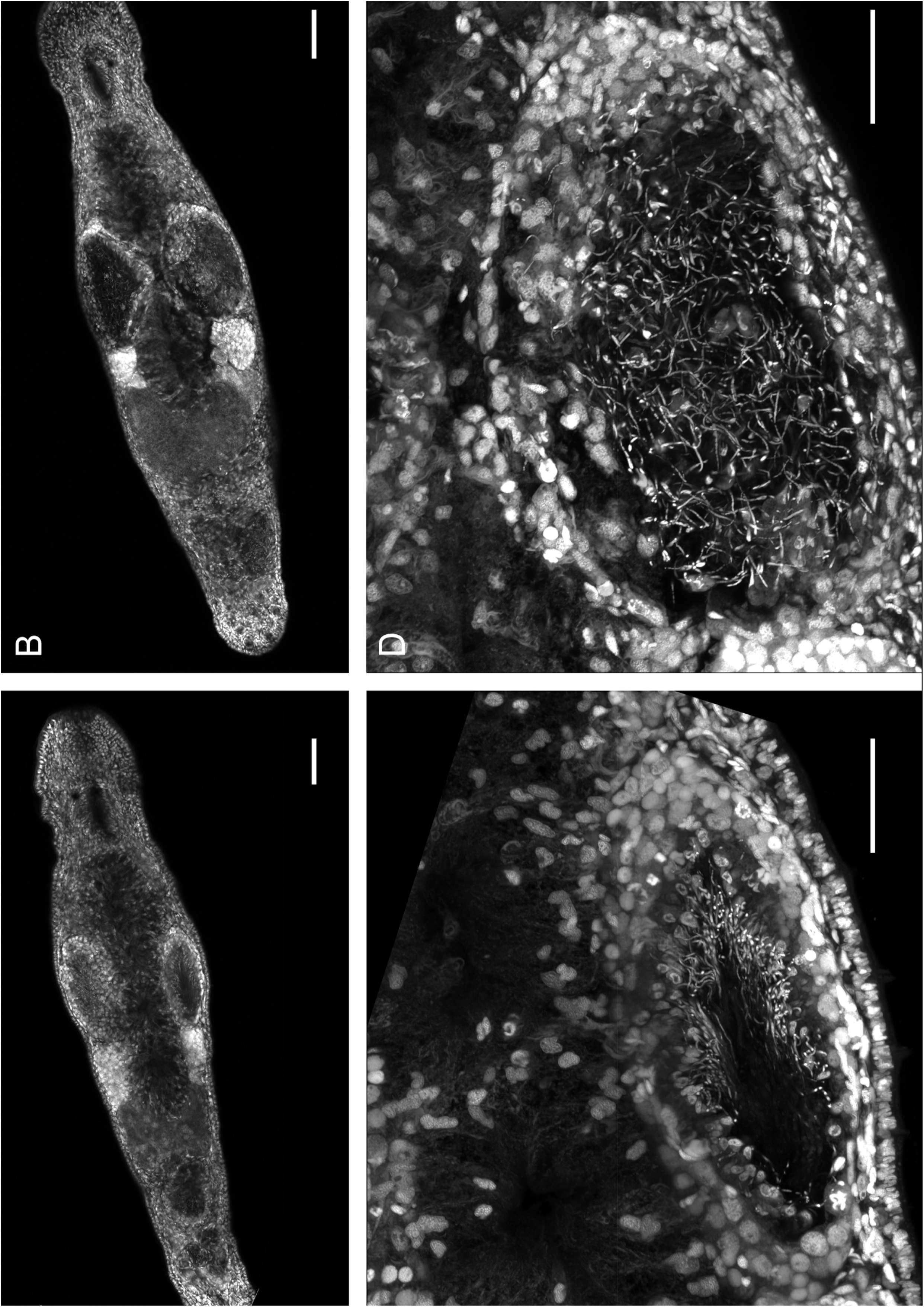
Single soaking with *sperm1* dsRNA is sufficient to obtain a compelling phenotype in *M. lignano*. Worms single-soaked with *sperm1* dsRNA or *gfp* dsRNA in regular f/2 medium (2 µg in 400 µL, 80 worms per assay) for 10 days, stained using the whole mount Carmin Red staining protocol, and imaged using confocal microscopy. Control worms soaked with *gfp* dsRNA display regular gonads and properly organized spermatozoa within the testis (A, C), whereas worms soaked with *sperm1* dsRNA display accumulated and strongly disorganized spermatozoa within the testes and swelling of the testes (B, D). Scalebars: 50 µm.

### Starving, absence of diatoms, and temperature are essential parameters for an optimal RNAi

The parameters of the single soak experiments in regular f/2 were based on the prerequisites of the electroporation method we used at the beginning of this study. Indeed, while the classical method used a daily renewal of the dsRNA in the presence of diatoms as a food source, our single soak approach is characterized by a short starving period of about 24 - 48 h before soaking and the absence of diatoms during dsRNA exposure. The presence/absence of diatoms during dsRNA soaking could thus be a critical parameter impacting the RNAi efficacy. Therefore, we conducted a comparative single soak experiment with *gfp* dsRNA on fed or starved NL1 worms in the presence or absence of diatoms. The GFP fluorescence in worm lysate was measured 12 days after the treatment for each condition (Fig 5A). Interestingly, these results suggest that both starving and the absence of diatoms in the f/2 medium strongly improve RNAi efficacy and reduce variability. It is worth noting that *M. lignano* worms puke out large amounts of digested diatoms after feeding. Therefore, a starving period followed by transferring the worms to a fresh medium lacking diatoms seems to be important to avoid the unintended presence of diatoms remnants.

**Figure 5:**
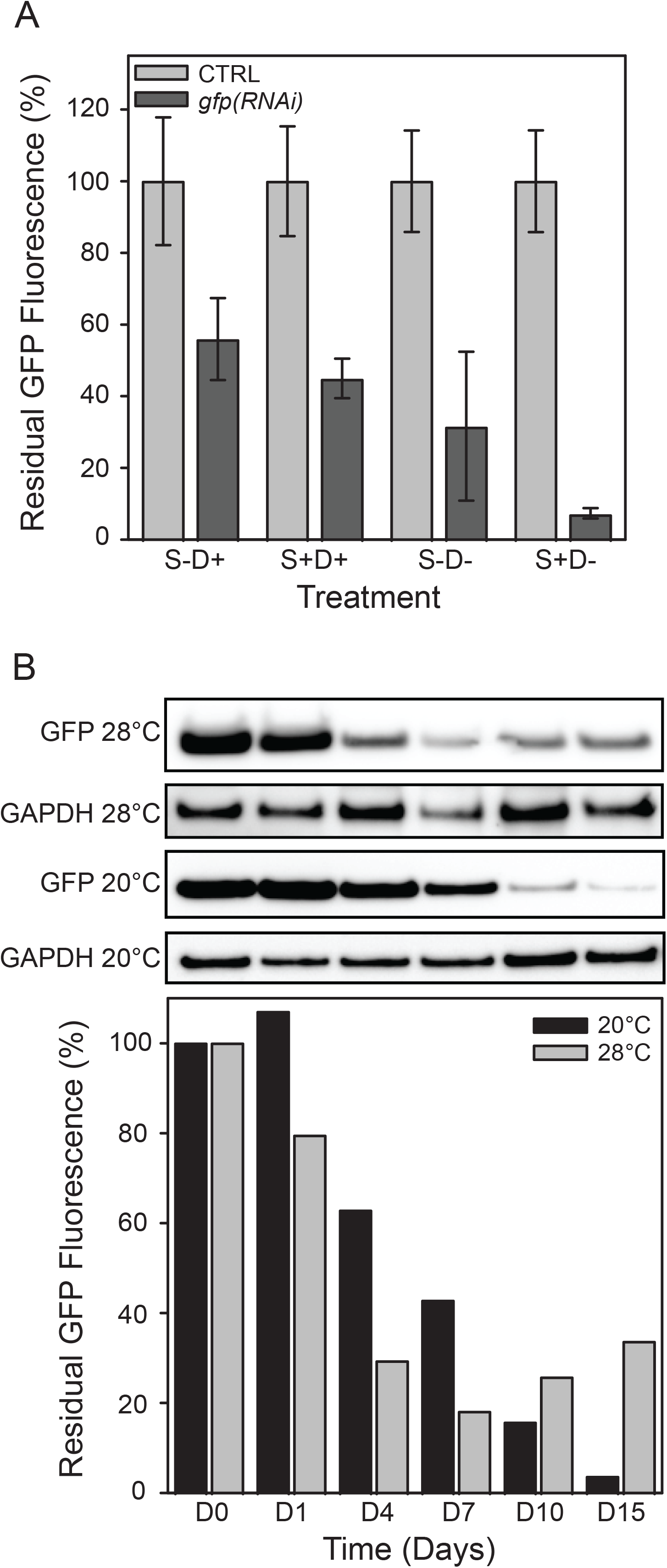
Starving, diatoms, and temperature are critical parameters for dsRNA efficacy. A. NL1 worms were starved or not for 24h and soaked with 2 µg *gfp* dsRNA (5 ng/µL) in the presence or absence of diatoms. Residual GFP fluorescence was measured (Ex=488 nm, Em=520 nm) 10 days after soaking and normalized to protein content as a percentage of fluorescence at D0 for each condition. B. Quantification of residual *gfp* expression after electroporation of 2 µg *gfp* dsRNA (5 ng/µL, 300 worms total) at 20°C and 28°C. For each time point up to 15 days, 30 worms were selected. GFP fluorescence was measured (Ex=488 nm, Em=520 nm) and normalized to protein content as a percentage of fluorescence at D0. For each time point, residual GFP expression was revealed by Western blot (16 µg total protein load) with an anti-GFP antibody (27 kDa). GAPDH is used as a loading control (36 kDa).

Since it was shown that increased temperature results in the faster manifestation of RNAi phenotypes in *M. lignano* (Wudarski *et al*. 2019), RNAi experiments have been performed at different temperatures between 20°C and 30°C. To test the influence of temperature on gene knockdown, we performed a quantitative comparison of RNAi efficiency at 20°C and 28°C. For this, a single soak in f/2 medium with dsRNA was performed, and worms were maintained at both temperatures. During a period of 15 days, the fluorescence intensity of GFP was measured together with Western blot analysis on a subset of worms at frequent time intervals (Fig. 5B). As expected, the residual GFP fluorescence decreased faster at 28°C than at 20°C, but interestingly, it also increased again earlier at 28°C than at 20°C. Moreover, the lowest residual GFP fluorescence was observed at 20°C, suggesting that the impact of RNAi is slower but potentially more efficient at 20°C.

### Production of dsRNA in bacteria is a high-yield and affordable method for RNAi

*In vitro* synthesis of dsRNA is the most expensive part of RNAi experiments in *M. lignano*. It requires multiple PCR steps and purification using cDNA templates synthesized directly from RNA or cloned into a plasmid. The dsRNA is generated by enzymatic transcription from linear DNA fragments initiated from two T7 promoters in the opposite orientation. We developed a more straightforward approach to extract dsRNA directly from bacteria containing cloning plasmids based on previous works (Nwokeoji *et al*. 2016; Sturm *et al*. 2018). While a regular *in vitro* synthesis using a plasmid template for PCR amplification results in ∼100 µg of purified dsRNA and requires two working days and relatively expensive reagents, the purification of dsRNA directly from bacteria yields >200 µg of dsRNA per 20 mL of culture, takes 2-3 working days, and relies on common laboratory reagents, substantially reducing the costs.

To compare the efficacy of bacteria-derived dsRNA with dsRNA that is enzymatically synthesized *in vitro*, we performed RNAi experiments targeting *ddx39b* and *piwi*. For both genes, the gene knockdown was similar and achieved >90% (Fig. 6A). To further evaluate the quality of bacteria-derived dsRNA, we studied its phenotypic effect in worms using the single soak approach in f/2 medium. Experiments resulted in clear regenerative phenotypes after tail amputation for *ddx39b*(*RNAi*) and *piwi*(*RNAi*) (Fig. 6B). By using the NL24 transgenic line, a clear decrease of fluorescence could be observed when *mNeonGreen*(*RNAi*) and *mScarlet*(*RNAi*) worms were compared to the *heh1*(*RNAi*) negative control worms (Fig. 6C). The *C. elegans heh-1* gene does not have a homolog in *M. lignano*, and can therefore be consistently used in all transgenic *M. lignano* lines expressing diverse fluorescent proteins. Altogether, bacteria-derived dsRNA is as effective as enzymatically synthesized dsRNA, but the production is cheaper and is easily scalable to generate large amounts of dsRNA.

**Figure 6:**
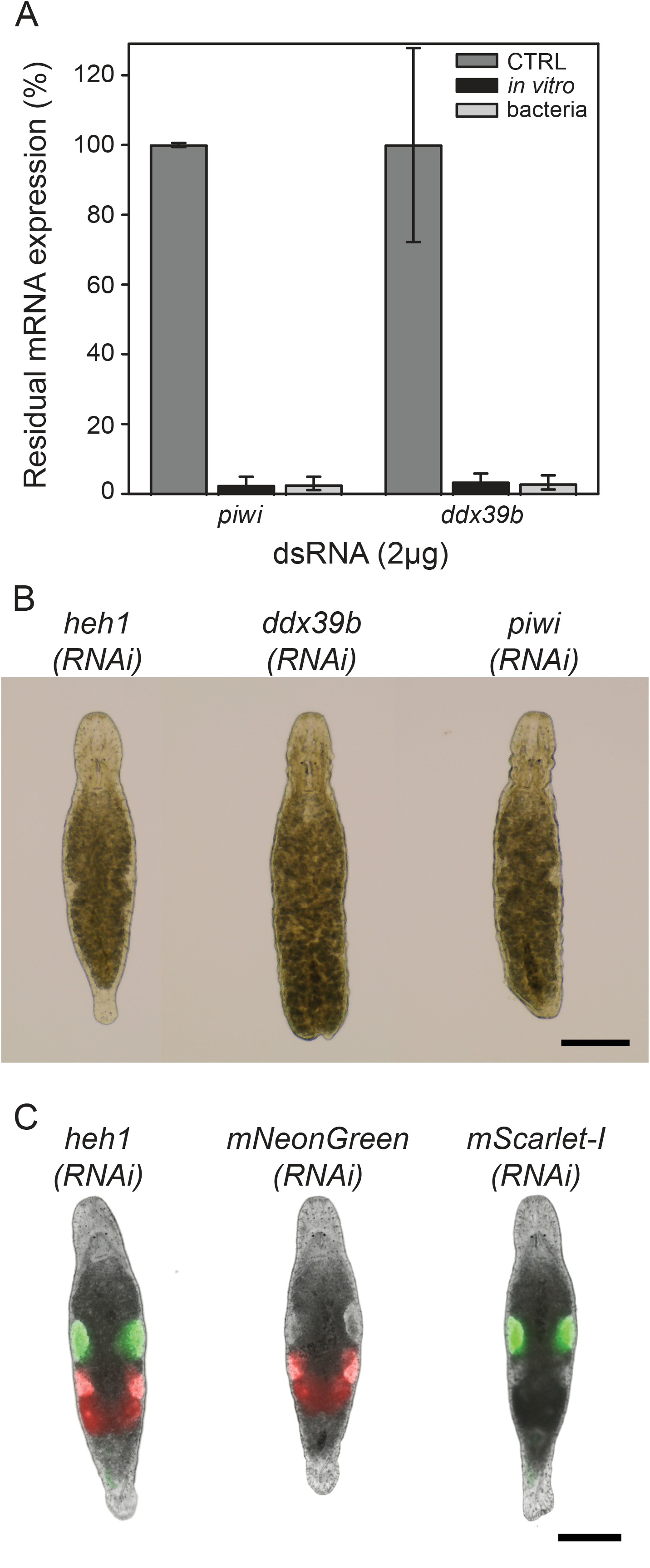
dsRNA produced in bacteria can drive efficient RNAi in *M. lignano*. A. Quantification of residual *piwi* or *ddx39b* mRNA expression using RT-qPCR (percentage of residual expression, +/- S.D.) 4 days after single soaking in regular f/2 with 2 µg of *piwi* or *ddx39b* dsRNA produced by *in vitro* synthesis or purified from bacteria (80 worms per assay, n=3, CTRL: no dsRNA). B. Regenerative phenotypes after single soaking in regular f/2 with 2 µg of *piwi* or *ddx39b* dsRNA produced in bacteria. The tail was amputated 5 days after the single soak and photos were taken 3 days after amputation. *heh1(RNAi)* represents the negative control (15 worms per condition, scalebar: 200 µm). C. Fluorescent phenotypes after single soaking of NL24 worms in regular f/2 with 2 µg of *mNeonGreen* (NG) or *mScarlet* (SC) dsRNA produced in bacteria. Photos are taken 7 days after the single soaking. Scalebar: 200 µm.

## Discussion

This study aimed to develop an optimized RNAi method for *M. lignano* with a reduced workload and a better cost efficiency while keeping the high efficacy of the traditional soaking approach. The conventional method, which is commonly used and validated in the community, is based on a daily renewal of the dsRNA solution in multi-well plates containing diatoms as a feeding substrate for worms. This daily renewal is factually time-consuming, uses a significant amount of dsRNA, and requires repetitive manipulations of worms, which, importantly, can induce worm stress and an increased risk of worm injuries and/or loss.

Inspired by our experience with RNAi in the blood-dwelling parasite *S. mansoni*, we first investigated the feasibility and efficacy of an electroporation method to introduce dsRNA in *M. lignano*. After adjustment of the “Schistosome-based” protocol to this marine organism, we demonstrated that electroporation effectively decreases gene expression and protein levels of target genes in a single step and results in clear, long-lasting phenotypes (Fig. 1). However, in analyzing the parameters driving the RNAi efficacy, we discovered that the electroporation protocol can be further simplified since the RNAi phenotypes were also observed in the control animals, where the electric pulse was omitted. At first, we suspected that the osmotic stress required to perform the electroporation in diluted f/2 medium was the main driver of dsRNA absorption by worms, but we established that regular f/2 medium was also sufficient to promote RNAi. This resulted in the most direct RNAi method: a single soak of 24 h or less in a regular f/2 medium with dsRNA at a concentration as low as 1 ng/µL. For many target genes, this method results in a long-lasting and efficient knockdown of gene expression and the development of phenotypes (Fig. 3, 4, 5B). As this approach requires very little hands-on time, this protocol will be best suited for larger RNAi screens and the preferred go-to approach for starting RNAi experiments in *M. lignano*.

From our previous experience, gene knockdowns by RNAi in *M. lignano* often do not result in obvious phenotypes. Once the effectiveness of the knockdown at the RNA level is established, but still no phenotype is observed, this often can be attributed to the functional redundancy and robustness of the biological process in which the gene is involved. This sometimes can be solved by additional sensitizing of the animals. For example, in the case of *tim29*(*RNAi*), a single soak was not always sufficient to develop a phenotype (Fig. 2D). This might be particularly true for studies of lowly-expressed genes or genes with very slowly developing phenotypes (Hong *et al*. 2014; ONCHURU AND KALTENPOTH 2019). There are several straightforward options to sensitize experiments. The first option is to repeat the single soak regularly, e.g., once a week. This enables long-lasting RNAi experiments with a potentially larger efficiency. Second, experiments can be sensitized by adding a stressor like osmotic stress, as demonstrated by the *tim29*(*RNAi*) example. Third, electroporation represents a powerful method to sensitize RNAi experiments as the small damages created all over the body cause a regenerative response without losing specific organs and tissues, as is the case with amputations. In addition, environmental parameters, such as temperature, can impact phenotype dynamics (Fig. 5B).

When dsRNA penetrates into cells, the interference of gene expression is almost immediate, while expected phenotypes could require days or weeks to appear, mainly because of protein half-life and delays in the subsequent biological responses. In order to ascertain RNAi efficacy, we combined quantitative biochemical analysis such as RT-qPCR, Western Blot, and fluorescence measurements (Fig. 3). Indeed, RT-qPCR allows a direct validation of proper dsRNA design and degradation of the targeted mRNA immediately after soaking, before associated phenotypes are observed or measured. Western blot analysis is complementary since it confirms the decrease of protein levels and also gives an indication of protein half-life and relative abundance; it is, however, limited by the availability of antibodies or biochemical tools. GFP fluorescence measurement is also a direct and convenient manner to precisely monitor RNAi efficacy and to analyze kinetics.

To give a more detailed description of phenotypes, we also adapted the Carmine Red staining protocol, originally used to analyze phenotypes on *S. mansoni*, for *M. lignano*. This staining protocol allows high-resolution imaging of whole-mount worms at the cellular level (Fig. 4). In addition, the Carmine Red staining enables the conservation of samples for years at room temperature without loss of fluorescence signal.

Often, dsRNA against the *gfp* gene is used as a negative control in RNAi experiments, but this is not appropriate for *gfp*-expressing transgenic lines. Therefore, we introduced *heh1* as a consistent target gene for the negative control condition in diverse transgenic lines to enable the design of robust, consistent RNAi experiments.

Independently of the protocol used, the highest cost of RNAi experiments performed in *M. lignano* remains the production of dsRNA. The classical soaking protocol requires a significant consumption of dsRNA over a 2 to 3 weeks experiment, *i.e.* up to 40 µg of dsRNA for a single well, traditionally containing 15-20 worms in 300 µL of f/2 medium. With our “one shot” soaking method, a maximum of 2 µg of dsRNA is required while treating up to 100 worms at a time. To decrease this cost further, we adopted an approach to produce and purify dsRNA directly from bacteria based on previously published studies (Nwokeoji *et al*. 2016; Sturm *et al*. 2018). The method is easy to perform and requires standard and widely available laboratory reagents and a simple microbiological setup. If necessary, the yield (>200 µg/20 mL of culture) is easily scalable by simply increasing the production culture volume. A new batch of dsRNA can be obtained within one day by re-inoculating a glycerol stock with the cloned plasmid. A universal cloning strategy through Gibson assembly simplifies the cloning procedure and screening of colonies and ensures the directionality of the insert. As a bonus, the same plasmid, universal and specific cloning and sequencing primers (Table S1) can be used to generate antisense ssRNA for *in situ* hybridization. Importantly, the protocol has already been successfully tested by an independent research laboratory (Erin Davies, NIH, personal communication).

During our studies, we wondered about the higher-than-expected efficiency of RNAi after a single soak in f/2 medium, especially compared to the published daily renewal of dsRNA solution. A crucial difference between our novel and the traditional soaking methods is the absence of diatoms during dsRNA exposure. Typically, soaking the worms in dsRNA solution is done in the presence of a pre-grown diatom layer to ensure food availability. In contrast, our optimized protocol only transfers worms into wells with pre-grown diatoms after the dsRNA exposure. In addition, worms are starved for 24 – 48 h before treatment to avoid the excretion of diatom remains in the presence of dsRNA. By limiting the total starvation period to a maximum of 48 h, its impact on the biology of worms can be minimized, while the presence of diatoms during dsRNA exposure is almost completely avoided. Our experiments confirm that this results in a more efficient and reproducible gene expression knockdown (Fig. 5A). The inhibitory effect of diatoms against RNAi efficacy by soaking can be explained by the chemical composition of the silicate diatom shell and the presence of a bacterial community living with these unicellular algae. Diatom shell is made of hydrated silicate and is an industrial source for the production of biochemical reagents to immobilize nucleic acids (Boom *et al*. 1990; Rea *et al*. 2014). Therefore, it is very likely that the diatom layer feeding the worms could limit the diffusion of the dsRNA into worms. Indeed, the mechanism promoting the entry and processing of dsRNA into *M. lignano* cells is still not described. In *C. elegans,* dsRNA molecules can be uptaken by specific transporters, SID-1 and SID-2, expressed in the gut epithelium (Winston *et al*. 2007; SHIH AND HUNTER 2011; Mcewan *et al*. 2012). Even if all the dsRNA import machinery and siRNA processing genes are present in *M. lignano* (Fontenla *et al*. 2021), the specific entry mechanism by the gut or the epidermis during dsRNA soaking has not been characterized so far. Therefore, one may argue that dsRNA binding to the diatom shell could disfavor its absorption through the gut. In addition, diatoms carry a complex microbiota hosted in the biofilm they produce (SISON-MANGUS *et al*. 2014). These bacteria could strongly contribute to the digestion of dsRNA (Bruckner *et al*. 2011) together with the adsorption/absorption by diatoms.

Overall, our experiments strongly suggest that a single soaking of *M. lignano* animals with dsRNA is sufficient to induce a strong and sustained RNAi when performed in the absence of diatoms. In addition, electroporation and osmotic shock are efficient sensitizers to promote RNAi when targeted genes are more problematic, while keeping a reduced workload and dsRNA production cost. These optimized and straightforward protocols represent a valuable resource for future RNAi experiments in *M. lignano* and other *Macrostomum* species. In addition, the developed protocol for dsRNA production in bacteria provides an efficient, cheaper method that can also be used in other model organisms.

## Acknowledgments

We warmly thank Mrs. Marie-José Ghoris for her technical support. We thank the laboratories of Dr. Erin L. Davies and Dr. Alexandr Blinov for independently testing the novel methods of dsRNA production in bacteria and single soak RNAi in *M. lignano*, and providing their feedback.

We thank Dr. Elizabeth Werkmeister and Sophie Salome-Desnoulez from the BiCel imaging facility in Lille for their precious support in confocal microscopy and image reconstitutions.

This work has been funded by the Dutch Research Council grant to EB (grant number OCENW.KLEIN.054) and by CNRS, INSERM and University of Lille recurring endowment (UMR9017-U1019) to JV.

## Notes

https://zenodo.org/records/10046394

### Competing Interest Statement

The authors have declared no competing interest.

