## Supplemental Material and Methods for "Optimized protocols for RNA interference in *Macrostomum lignano*"

### Double-stranded RNA (dsRNA) production in HT115 *E. coli* cells from T444T-based plasmids and dsRNA purification

#### Part I. Cloning of a part of a target gene of interest (GOI) (3 days of hands-on work)

***Day 0: Preparation.***
1) Design and order a pair of primers to amplify ~400-600 bp of GOI from either gDNA (the highest chance of success) or cDNA. Add **ATAGGGAGACCGGCAGAT** 5' overhang to the forward (FWD) primer, and **ACTATAGGGCGAATTGGGTA** 5' overhang to the reverse (REV) primer.

***Day 1: PCR amplification and cloning of GOI.***
2) Amplify the selected sequence region by PCR using a proofreading DNA polymerase (like Q5 (NEB) or KAPA (Roche)).

3) Gel-purify the PCR product(s) either by a 'Freeze&Squeeze'* or spin-column purification method.
***Note:*** *Freeze&Squeeze is a fast and cheap in-house method for small and large-scale gel extractions. Briefly, excised gel blocks are frozen in 1.5 mL tubes at -20°C (1 h) or -80°C (15 min), or liquid nitrogen (immediately). Then, the tubes are centrifuged at the maximum speed at room temperature for >3 min to crack and squeeze the gel. Gel-free supernatants are suitable for common subsequent molecular biology applications: ligation, restriction digestion, PCR, Gibson Assembly, Sanger sequencing, etc. The presence of the gel-loading dye does not affect the subsequent reactions.*

4) Using the NEBuilder® HiFi DNA Assembly Cloning Kit (NEB) or another 2xGibson assembly MasterMix, mix on ice the following components in a 200 µL PCR tube for one reaction:

| **Reagent** | **Volume (µL)** |
|---|---|
| 2xNEBuilder HiFi MasterMix | 1 |
| BglII and KpnI digested and gel-purified T444T plasmid* | 0.5 |
| The gel-purified PCR product | 0.5 |

** prepare a sufficient amount of the cut and gel-purified T444T vector (Sturm et al. 2018) in advance and store it at -20 °C. Use of the BglII and KpnI restriction enzymes is mandatory if you will use the 5' primer overhangs from this protocol.*

Put a drop of a mineral/halocarbon oil to cover the reaction mix completely. Briefly centrifuge the tube if necessary. Incubate the reaction at 50°C in a thermocycler for 30-60 min. Afterwards, transfer 2 µL of the reaction to a 1.5 mL tube.

5) Transform the NEBuilder reaction to competent **HT115** *E. coli* cells. Plate the cells on **LB + AMP (200 µg/mL) + TET (12.5 µg/mL)** agar. ***Note:*** *tetracycline (TET) can be added on top of the LB+Amp plates 1 h prior to the plating. To do it, add 10 µL of 1000x TET to 990 µL of LB and spread 150 µL of the mix on the agar plate.*

***Day 2: Checking the insertions***
6) [Optional] On the next day, check 2-4 colonies for insertions by colony PCR using either the Universal_T444T_T7_FWD (**TTATGCTAGTAATACGACTCACTATAGGG**) primer or a pair of the T444T_FWD_Seq2 (**CGGTTCCTGGCCTTTTGC**) and T444T_REV_Seq2 (**CCCGATTTAGAGCTTGACG**) primers.

Pick a colony using a 10 µL pipette tip and mix the colony in 15 µL milliQ $H_2O$ in a 200 µL PCR tube. Attach the same tip to a P2 or P10 pipette and transfer 1 µL of the colony mix to a new PCR tube. Store the colony mixes at 4°C. Prepare a master mix using OneTaq® Quick-Load® 2X Master Mix with Standard Buffer (NEB):

| **Reagent** | **Volume (µL)** | **Master Mix (µL)** |
|---|---|---|
| 2xOneTaq Mix | 5 | $5*(N_{colonies}+1)$ |
| Universal_T444T_T7_FWD | 1 | $N_{colonies}+1$ |
| $H_2O$ (milliQ) | 3 | $3*(N_{colonies}+1)$ |

Add 9 µL of the PCR Master Mix to 1 µL of the colony mix. **Tip: do reverse pipetting to avoid formation of bubbles.**

Run the following thermocycler program:

| | | |
|---|---|---|
| 94°C | 5 min | 1 cycle |
| 94°C<br>58°C<br>68°C | 10 s<br>10 s<br>45 s | 20 cycles |
| 68°C | 1 min | 1 cycle |
| STOP the program | | |

Check the PCR products on 1% agarose gel stained with Ethidium Bromide.

7) Select 1 positive colony per construct. Add the other 14 µL of the colony mix to 4-5 mL of liquid **LB + AMP (200 µg/mL) + TET (12.5 µg/mL)**. Grow at 37°C overnight with shaking at 180-200 rpm.
***Note:*** *Tetracycline is only necessary to ensure specific growth of the HT115 E. coli strain relative to other lab strains that can have Amp resistance. However, you can omit TET here and at the subsequent steps, if there are no doubts of potential cross-strain contamination.*

***Day 3: Plasmid isolation, glycerol stock preparation, and sequencing:***
8) The next morning, take 0.5 mL of the overnight culture and mix it with 0.5 mL of 50% autoclaved glycerol in milliQ. Work sterile. Freeze the glycerol stock in liquid nitrogen and/or put it directly to -80°C.

9) From the rest of the overnight culture, isolate the plasmid by miniPrep or any other plasmid isolation protocol.

10) Send the plasmid for Sanger sequencing with either of the T444T sequencing primers: T444T_FWD_Seq2 (**CGGTTCCTGGCCTTTTGC**) or T444T_REV_Seq2 (**CCCGATTTAGAGCTTGACG**).
***Note:*** *due to the cloning method the orientation of the insert is always known and is the same as of the FWD primer.*

11) Store the rest of the plasmid at -20°C as a backup in case of failed glycerol stock preparation.

#### Part II. dsRNA production in HT115 *E. coli* cells (1 day of hands-on work)

1) Take a glycerol stock of HT115 *E. coli* cells with the GOI from the -80°C freezer. Do not defrost the cells. Work near the flame. Using a flame-sterilized microbiological loop, scrape for 2-3 seconds at the top of the frozen glycerol stock and inoculate 4 mL of fresh liquid **LB+AMP(+TET** [Optional]**)** medium.

2) Grow the cells overnight at 37°C with shaking at 180-200 rpm.

3) The next morning, pellet the cells, remove all the supernatant, and resuspend the pellet in a fresh LB+Amp medium. Add 200 µL of the overnight culture to 20 ml of fresh liquid **LB + 200 µg/ml AMP (NO TET!)** medium in a 50 mL falcon tube. Close the tube with aluminum foil instead of the lid to enable airflow, but keep the lids for later.
***Notes:*** *Important! (1) Dilute ampicillin from the stock on the same day as culture growth. Do not use old LB+Amp. The concentration of Amp here is higher than the standard 50-100 µg/mL to prevent loss of the plasmid by the bacteria. (2) The volume of the production culture can be scaled up. Increase the amount of the inoculating overnight culture proportionally. If you will work with glassware, autoclave it filled with milliQ water to eliminate residual detergents that can interfere with bacterial growth.*

4) Grow the culture for 3 h at 37°C with shaking (at 180-200 rpm) to reach the mid-exponential growth phase. Afterwards, add 100 µL of 0.1 M IPTG. Continue to grow the culture for an additional 4-5 h.
***Note:*** *if the volume of the production culture was scaled up. Increase the amount of IPTG proportionally.*

5) Upon the growth completion, close the tube culture with the saved lids and pellet the cells at 4000 rpm for 5-10 min. Remove the supernatant completely and freeze the pellet by storing it at -20°C until you are ready for RNA extraction.
***Note:*** *if the volume of the production culture was scaled up, split the culture into several 50 mL tubes for centrifugation.*

#### Part III. Nucleic acid isolation from the *E. coli* cell pellets (1 day of hands-on work)

*This step is a modified version of the RNASwift dsRNA extraction protocol (Nwokeoji et al. 2016, Nwokeoji et al. 2017)*

***Step 1: nucleic acid extraction***
Be sure to have prepared the following buffers in advance:

1) The Lysis Buffer: (4% SDS, 0.5M NaCl in milliQ $H_2O$)
2) 5M NaCl in milliQ $H_2O$;
3) 70% EtOH in milliQ.
4) 100% Isopropanol.
5) 1x DNAse I buffer (100 mM Tris/HCl, 25 mM $MgCl_2$, 1 mM $CaCl_2$ in milliQ)

1) Briefly preheat the Lysis Buffer buffer in a microwave to dissolve SDS and keep it warm (60-80°C).
**Note:** *if the volume of the growth culture was higher than 20 ml, adjust the volumes of the buffers proportionally (see below).*

2) Add 665 µL of the warm Lysis Buffer to the defrosted cell pellet and resuspend cells by pipetting. Immediately transfer the mix to a 2 mL tube and incubate it at 80°C for 10 min. This should result in a much clearer lysate mixture. Take the tube out and leave it at room temperature for 10 min to gradually cool down.

3) Add 335 µL of 5M NaCl and mix by shaking/inverting the tube to ensure sufficient mixing. This should result in a white milky precipitation of SDS.

4) Centrifuge the tube at the maximum speed (>18000 xG) for 10 minutes or longer. Transfer the supernatant to a new 2 mL tube. Try to minimize carry-over of the SDS pellet.
**Note 1:** *If a dense foamy film is formed at the top of the supernatant, remove it mechanically by picking with a p10/p200 pipette tip.*
**Note 2:** *If possible, centrifuge at 4°C to prevent non-specific RNA degradation.*

5) Add 950 µL (1 volume) of 100% isopropanol and mix well by vigorously shaking/inverting or by pipetting with a P1000 tip.

6) Centrifuge the tube at the maximum possible speed for 15 min. Decant the liquid (be careful not to disturb the pellet) from the tube.

7) Add 1 mL of 70-75% EtOH and invert the tube several times to wash its walls. Centrifuge at max speed for 5 min. Decant the liquid supernatant.

8) Repeat the wash step one more time.

9) Aspirate as much liquid as possible with a P10-P20 pipette. Air-dry the pellet for 10-15 min.

10) Add 500 µl of warm (~40-50°C) 1x DNAse I buffer and resuspend the pellet. Transfer the mix to a 1.5 mL Eppendorf tube.

11) Incubate the tube at room temperature or 45-50°C (recommended) for 5-10 min until the pellet is completely dissolved. Add more solvent if the pellet is still not dissolved.

12) Add 2 µL of DNAse I and 2 µL of RNAse T1 and incubate for 30 min at 37°C to eliminate most of the residual *E. coli* gDNA, ssRNA, and plasmid DNA. Mix by pipetting or inversions (do not vortex!)
**Note:** *Some E. coli rRNA will still be present due to its secondary structure. If it is not possible to use RNAse T1 and DNAse I, you can attempt purification by LiCl precipitation described as a step in the manual of the Replicator RNAi Kit (http://www.ulab360.com/files/prod/manuals/201305/13/78762001.pdf).*

***Step 2: removal of the E. coli 5S dsRNA fraction (~150 bp) and other small nucleic acids (PEG-salt size selection)****
**** an alternative, although a more expensive but much faster method, would be to use SPRI magnetic beads like AMPure DNA XP or SPRIselect (Beckman Coulter)***
Be sure to prepare the following buffers in advance:

1) 50% PEG8000 in miliQ $H_2O$ (w/v);

2) 5M NaCl in miliQ $H_2O$;
3) 70% EtOH in miliQ/RNAse-free $H_2O$

***Note:*** *all the volumes below are calculated for 500 µL of dsRNA solution. The final concentrations are 11% PEG8000 and 0.5 M NaCl. The cut-off value can be adjusted by varying PEG concentrations in the solution (less PEG – higher cut-off (>= 150 bp)).*

12) Centrifuge the mix at full speed for 5-10 min to pellet any residual SDS flakes. Gently and accurately transfer the dsRNA solution to a new 1.5 mL Eppendorf tube and add 74 µL of 5M NaCl and 166 µL of 50% PEG8000. Mix everything well by inverting the tube until homogeneous.

***Important note 1:*** *Residual SDS flakes can kill the worms, and therefore any SDS carryover must be avoided.*

***Important note 2:*** *50% PEG is very viscous. Use a cut P1000 tip to slowly and accurately take up the required volume.*

13) Let the mixture sit at room temperature for 15-20 min. Then, centrifuge the sample at maximum speed at room temperature for at least 20 min.

14) Using a P200 pipette accurately aspirates the supernatant, paying attention not to disturb the pellet, which may be hard to see. Leave some supernatant behind if you are not sure.

15) Add 500 µL of 70% EtOH to wash away the salt and PEG. Centrifuge for 5 min at full speed. Carefully aspirate the supernatant and repeat the washing step.

16) Using a P10 pipette, aspirate as much of the supernatant as possible. Leave the sample to dry for 10 min at room temperature.

17) Resuspend the sample in 500 µL of nuclease-free water.

***Step 3: agarose gel dsRNA integrity analysis and approximate dsRNA concentration estimation***

18) Dilute 1 µL of the sample in 9 µL milliQ water. Add 2 µL of a 6x Gel Loading Dye (no EtBr added directly in the dye).

19) Prepare 1% TAE-based agarose gel with EtBr. Use a comb with wider wells. Mind leaving one well for the DNA molecular weight marker.

20) Depending on the expected concentration of your sample, load 10 µL of the DNA ladder like NEB 1kb DNA ladder (N3232L). Load all the prepared samples.

21) Run the gel at 80 V until there is good separation between the DNA ladder bands. After the run, take a photo of the gel.

22) If dsRNA profile is good, measure its concentration by spectrophotometry at 260 nm using NanoDrop or an equivalent set on RNA parameters.

23) Aliquot dsRNA in 15-20 µL samples (to reduce freeze/thawed cycles) and store at -80°C for several months.
