## Supplemental Table S1 for "Optimized protocols for RNA interference in *Macrostomum lignano*"

### Supplementary Table S1

| Primers | Forward (5'-3') | Reverse (5'-3') |
|---|---|---|
| ***Primers for Topo-PCR-4-TA and pGEM-T cloning systems*** | | |
| *Mlig-ddx39b* | GGGTGAACATCAGCGTGTAC | TCCTCTTCTCGAAGTTCTTG |
| *Mlig-sperm1* | TAGCCCTGCTAAGGCAGGTA | CGATGTTGGCGTTTCTTCTT |
| *Mlig-piwi* | TGCTCAAGCTGGTGTTGC | GTCTTGTTGTTGTGCCGC |
| *gfp* | CGTAAACGGCCACAAGTTCAG | GAACTCCAGCAGGACCATGTG |
| *Mlig-tim29* | AAAGTTGAAAGCAAATTTCTGG | GCCAATCGATATAAAATTGGAA |
| pGEM-T | CGGCCGCCATGGCCGCGGGA | TGCAGGCGGCCGCACTAGTG |
| pGEM-T with T7 extension | ggatcctaatacgactcactatagg CGGCCGCCATGGCCGCGGGA | ggatcctaatacgactcactatagg TGCAGGCGGCCGCACTAGTG |
| ***Primers for in vitro dsRNA synthesis. T7 promotors are in lower case*** | | |
| *piwi* T7 | taatacgactcactatagggaga TGCTCAAGCTGGTGTTGC | taatacgactcactatagggaga GTCTTGTTGTTGTGCCGC |
| *ddx39b* T7 | taatacgactcactatagggaga GGGTGAACATCAGCGTGTAC | taatacgactcactatagggaga TCCTCTTCTCGAAGTTCTTG |
| *sperm1* T7 | taatacgactcactatagggaga TAGCCCTGCTAAGGCAGGTA | taatacgactcactatagggaga CGATGTTGGCGTTTCTTCTT |
| *gfp* T7 | taatacgactcactatagggaga CGTAAAC | taatacgactcactatagggaga GAACTCC |
| ***Primers used to PCR amplify fragments to clone into T444T vector for bacterial production of dsRNA. Overhangs (oh) to T444T are in lowercase*** | | |
| *ddx39b*-ohT444T | actatagggcgaattgggta TGCATGAACTTCTTGCAGATTG | actatagggcgaattgggta TGCATGAACTTCTTGCAGATTG |
| *mNG*-ohT444T | actatagggcgaattgggta ACGTCGGTGAAGGCCTTCT | actatagggcgaattgggta ACGTCGGTGAAGGCCTTCT |
| *mSC*-ohT444T | actatagggcgaattgggta ACGGTGTAGTCCTCGTTGT | actatagggcgaattgggta ACGGTGTAGTCCTCGTTGT |
| *piwi*-ohT444T | actatagggcgaattgggta GTCGCCGTTCTTCTCTCTGA | actatagggcgaattgggta GTCGCCGTTCTTCTCTCTGA |
| *heh1*-ohT444T | actatagggcgaattgggta CTGCTGGATGATTCTCGGTGA | actatagggcgaattgggta CTGCTGGATGATTCTCGGTGA |
| ***Primers for qPCR*** | | |
| *ribosomal S19* | GTCTAACCAATGGCTGACGACC | TCGGAGAACTGATCGATGCTCATG |
| *piwi* | AGGCCATTGTGGTGAAGAAG | ACTGCGACACCAGGAAGAAG |
| *sperm1* | GGCCGGCGGGGTGTATGA | AGTTGGCTGCAGCTCCGTCTCC |
| *ddx39b* | TTCACGGACTCAAGCAGCACTACCT | CCAGAACGTCGATCAGCTCAAACAG |
