## Supplemental Figure Legends for "Optimized protocols for RNA interference in *Macrostomum lignano*"

### Supplementary Legends

**Supplementary Figure 1:**
**Carmin Red staining in *M. lignano***
A. Reconstitution of the testes and ovaries of wild type *M. lignano* using projection of multiple plans of confocal images (Airyscan2 processing, maximum intensity projection mode) on whole mounted worms after Carmin Red staining. Scalebar: 50 µm.
B. Reconstitution of wild type *M. lignano* complete reproduction organs using 12 confocal images (Airyscan2 processing, stitching mode) on whole mounted worms after Carmin Red staining. Scalebar: 20 µm.

**Supplementary Video 1:**
3D reconstruction of *M. lignano* testis using multiple plans of confocal images on whole mounted worms after Carmin Red staining (Airyscan2 processing, zoom 1.8, 68 slices, deltaZ=0.5µm, pixel size=43 nm, tilted with progressive elimination plan, scalebar: 10 µm).

**Supplementary Video 2:**
3D reconstruction of *M. lignano* testis and spermatozoa using multiple plans of Airyscan2 processed confocal images on whole mounted worms after Carmin Red staining (Airyscan2 processing, zoom 1.8, 9 slices, deltaZ=0.5µm, pixel size=47 nm, zoom in and out tilted motion, scalebars: from 5 to 10 µm).
