## Supplementary figures and images for "Optimized protocols for RNA interference in *Macrostomum lignano*"

### Supplemental Fig 1

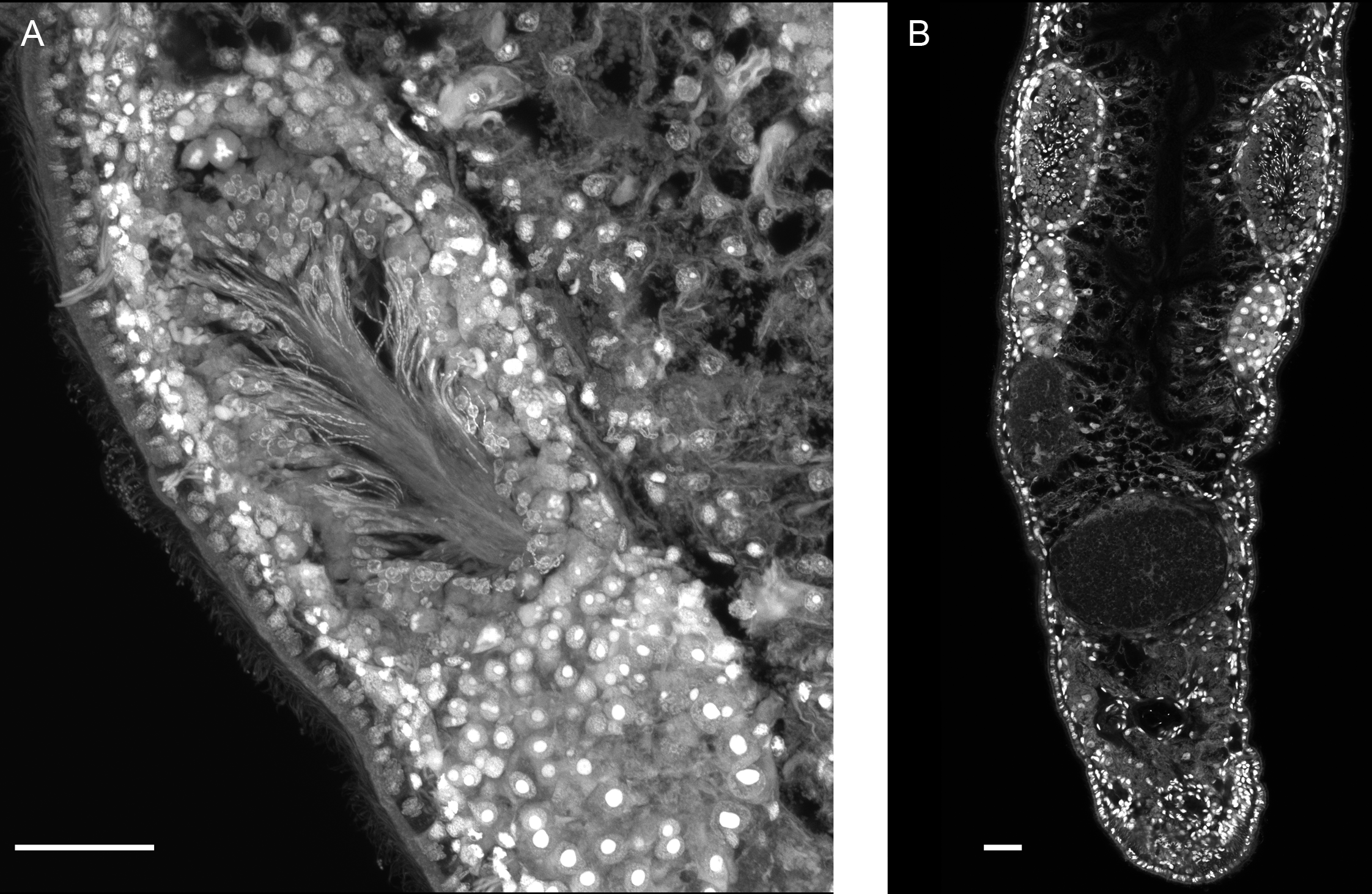
